# Microbiome Prevents Sudden Death Through Occult Cardiac Sub-micrometastasis in Mice

**DOI:** 10.1101/2023.07.03.547566

**Authors:** Syed Mohammed Musheer Aalam, Xiaojia Tang, PS Hari, Salem M. Salem, Ahmad Al Jarrad, Peter A. Noseworthy, Atta Behfar, Anguraj Sadanandam, Krishna Kalari, Purna C. Kashyap, Nagarajan Kannan

**Author notes:** Corresponding author: Nagarajan Kannan Ph.D., Director Stem Cell and Cancer Biology Laboratory, Mayo Clinic, 200 First Street SW, Rochester, MN, 55905, USA, Electronic mail.

## Abstract

Sudden cardiac deaths (SCDs) pose a formidable clinical challenge, and their underlying risk mechanisms are poorly understood. Using a gnotobiotic, germ-free mouse model, we fortuitously discovered SCD incidences resulting from occult cardiac metastases. Female germ-free C57BL/6 mice (n=22) were raised in isolation and injected with mammary Py230 cells. Significantly higher SCD probabilities (36.3%) were observed in germ-free mice compared to gut-colonized groups (0%). Extensive examinations revealed no physical anomalies but demonstrated occult cardiac sub-micrometastasis in three out of four sudden death cases. Further analysis supported the role of occult cardiac sub-micrometastasis as the leading cause of SCDs. The remaining germ-free mice exhibited minimal primary tumors but high levels of cardiac metastases and morbidity. The gnotobiotic SCD model represents a crucial milestone in our understanding of the complex interplay between the gut microbiota and the development of occult oncological processes that ultimately culminate in SCDs and warrants further investigation into their mechanisms.

## Main

In the United States, sudden cardiac deaths (SCDs) constitute a substantial proportion (5.6% to 15%) of all deaths, and approximately 50% of individuals experiencing SCDs have no prior diagnosis of a heart disorder ^1-4^. The lack of comprehensive understanding regarding the underlying causes of SCDs presents a formidable clinical challenge, given their unexpected nature and the limited efficacy of current preventive measures. Consequently, there is an urgent need to enhance our knowledge and unravel the etiological factors contributing to SCDs, with the ultimate goal of developing effective preventive strategies. By happenstance, we have discovered SCD incidences resulting from previously unknown microbially preventable occult cardiac sub-micrometastasis, whose clinically inscrutable origin is experimentally defined, in a novel germ-free mouse model.

To ensure the isolation of the microbiota-free environment, we raised female gnotobiotic germ-free C57BL/6 mice (n=22) in a germ-free isolator for the entire duration of the study. Moreover, we conducted experimental intragastric fecal microbiota transplantation on a batch of 5-week-old female germ-free (conventionalized; n=20) mice using a healthy mouse donor and included a group of mice naturally gut-colonized (conventional; n=10) at birth. At week 8, all three groups of mice were administered with subcutaneous injection of syngenic MMTV-PyMT oncogene-encoded and DNA barcoded multipotent mammary Py230 cells^5^ (experimental design shown in **Figure-1A**). The germ-free status was confirmed by colonic bacterial 16S PCR (**Figure-1B**)^6^.

**Figure 1:**
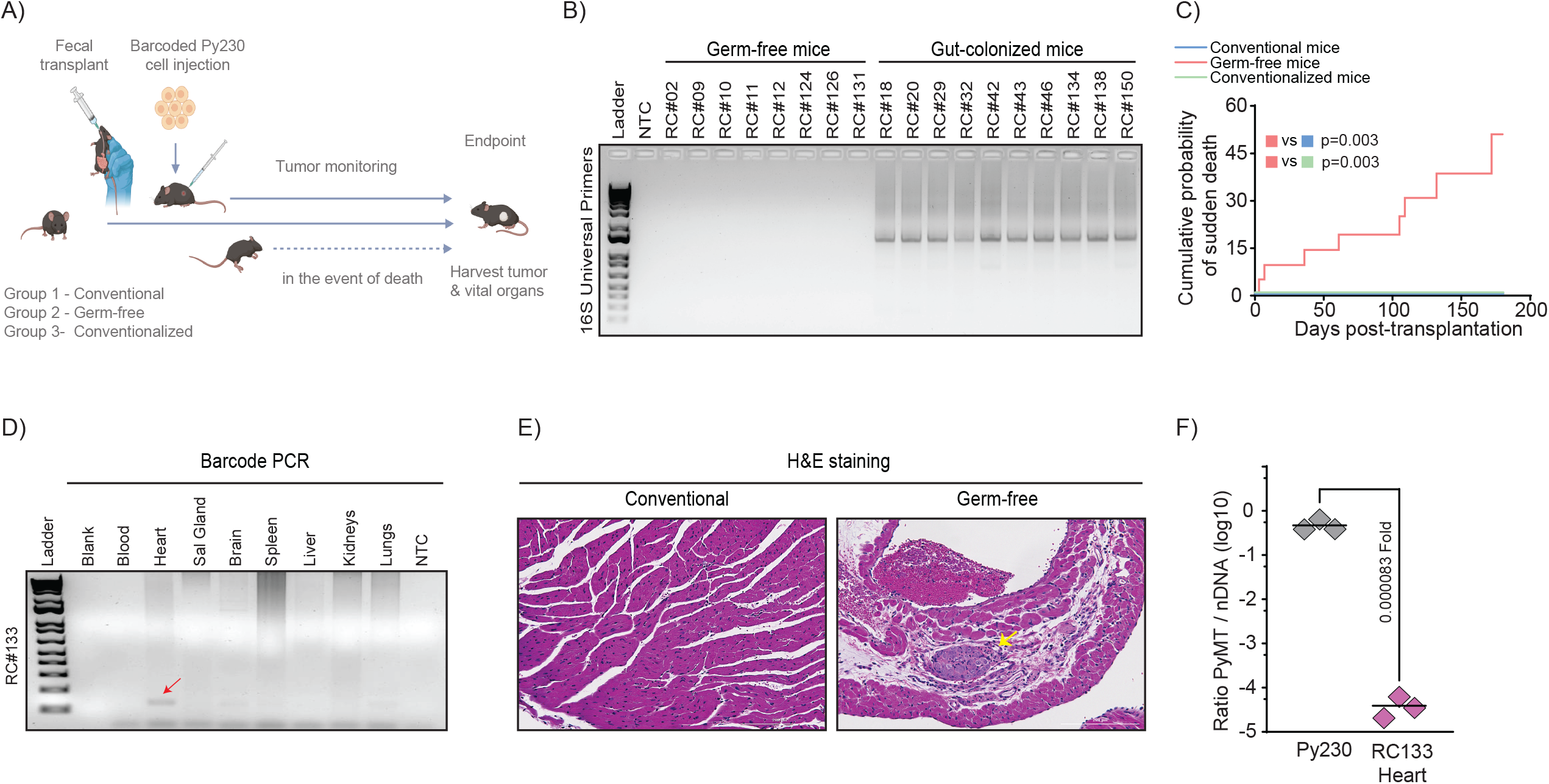
**[A]** illustration showing the experimental design of the study. **[B]** A representative gel electrophoresis image showing PCR amplification of the universal 16S rRNA genomic region in DNA extracted from colonic fecal samples collected from gut-colonized and germ-free mice. NTC is ‘no template control’ (NTC), and RC# is an individual animal identification number. **[C]** Kaplan-Meier plot showing the cumulative risk of SD during the first 6 months in the germ-free group compared to both conventional and conventionalized groups. The log-rank (Mantel-Cox) test was used for comparison of curves (p-values are indicated). Eight germ-free mice experienced SD. **[D]** A representative gel electrophoresis image showing PCR amplification of barcode sequences in the heart (indicated by a red arrow) but not in other vital organs of the SD germ-free mouse. NTC is ‘no template control’ (NTC). **[E]** Photomicrograph showing representative hematoxylin and eosin-stained images (x200 magnification) of heart tissue sections obtained from control and germ-free mice. A micro-metastatic lesion involving myocardium is indicated by a yellow arrow. **[F]** ddPCR plot showing the rarity of Py230 cells in the heart of a germ-free mouse (RC133) that experienced SD. The plot depicts the ratio of the transgenic PyMT gene (1 copy per cell) to the 18S rRNA gene (2 copies per cell) in the nuclear DNA (nDNA) obtained from parental Py230 cell line and heart.

There were significant differences observed in the probability of sudden deaths (SDs) in germ-free group compared to gut-colonized groups (conventional and conventionalized; p<0.005) (**Figure-1C**). The incidence of SDs was approximately 36.3% (8 out of 22 mice) in the germ-free group within 6 months post Py230 injection, compared to 0% in the conventional mice. The SDs were fully eliminated (0%) in germ-free mice upon colonization with gut microbes, as depicted in **Figure-1C**, highlighting the safeguarding effect of certain gut microbes against SDs in this mouse model.

To determine the cause of SD in eight germ-free mice, we examined their external features, such as body size, fur condition, and integumentary abnormalities, as well as their internal organs, with a focus on identifying inflammation or tumors. However, we did not observe any physical anomalies or neoplastic growth, which left us perplexed about the underlying pathophysiology responsible for the SD. DNA was isolated from four out of eight germ-free mice that experienced SD. The PCR analysis of Py230 cell markers, which were labeled with lentivirally marked DNA barcodes^7^, was performed on the vital organs of four mice that had experienced SD. The results indicated occult cardiac metastasis in three out of four SD mice (**Figure-1D**).

To gather further evidence of a cardiac event as the cause of SD, we conducted the following experiments: i) microscopic examination of hematoxylin and eosin-stained sections of myocardial tissues in mice from all three groups (**Figure-1E**); ii) droplet digital polymerase chain reaction (ddPCR) analysis in heart tissues of three SD germ-free mice, targeting the PyMT gene, which is a transgene encoded in the Py230 cell genome (**Figure-1F**); iii) Illumina HiSeq4000 platform-based analysis of DNA barcode sequences amplified by PCR from heart tissues of three SD germ-free mice, to investigate whether the occult cardiac metastasis was a result of rare monoclonal or oligoclonal metastatic event (**Figure-2A**).

**Figure 2:**
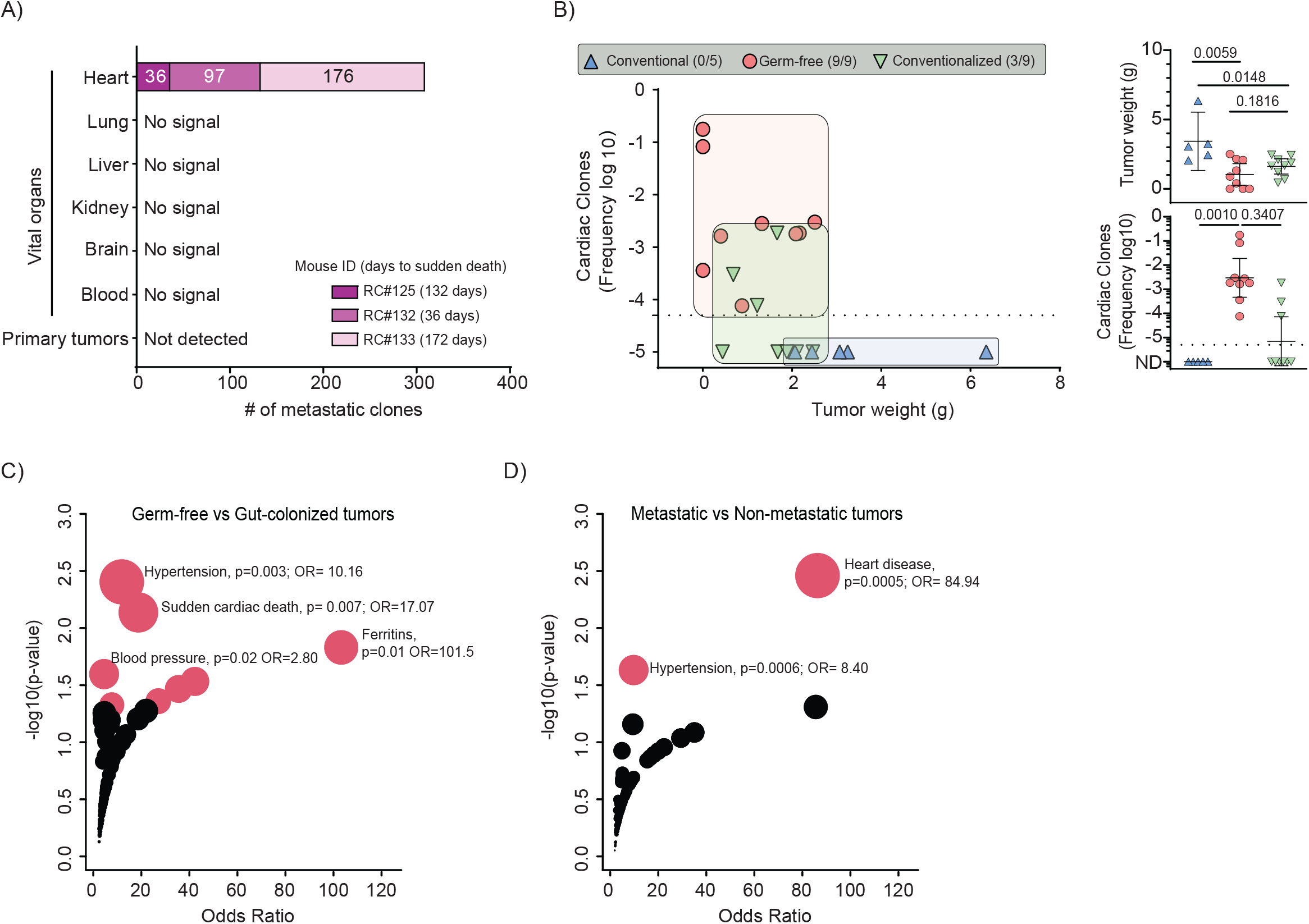
**[A]** Plot shows heart specific polyclonal enigmetastatsis in three SD mice (RC125, RC132, and RC133). **[B]** Plot showing no correlation between tumor weight and cardiac clonal frequencies in three groups of mice. Group-wise comparisons and p-values are indicated. **[C]** A plot showing various phenotypes associated with the DE genes in germ-free (n=7) vs gut-colonized (n=12) mouse tumors. **[D]** Similar plot showing phenotypes associated with DE genes obtained from metastatic (n=12) vs non-metastatic (n=7) tumors.

Microscopic examination revealed a rare cardiac sub-micrometastatic lesion in a germ-free mouse (**Figure-1E**). By analyzing up to three micrograms of DNA in multiple ddPCR analyses, we detected Py230 cells (1 in ∼800K cells) in the heart in one out of three SD mice examined. DNA barcode analysis identified the presence of polyclonal metastasis exclusively in the hearts of all three mice that suffered from SD (**Figure-2A**). The data generated herein provide strong evidence supporting the notion that occult cardiac metastasis is the most plausible cause of SD. The findings from our experiments, including the microscopic examination of myocardial tissue and the genetic analysis of Py230 cells, provide convincing support for this hypothesis.

Compared to conventional mice, the germ-free mice displayed small or no primary tumors (p=0.005), yet consistently higher levels of cardiac metastasis (p=0.001), indicating that occult and early cardiac metastatic events may be a prevalent characteristic in this model (**Figure-2B**). The link to cardiac events was further uncovered by comparing differentially expressed genes in tumors obtained from germ-free mice (n=7) with those from gut-colonized (conventional, n=4; conventionalized, n=8) mice that are resistant to similar events. Analysis of differentially expressed genes of primary tumors using the NCBI’s database of Genotypes and Phenotypes (dbGaP) of 12 metastatic and 7 benign tumor-bearing mice or 7 germ-free and 12 gut-colonized mice showed heart disease (p=0.0005/OR=84.94) and SCD as a significant phenotype (p=0.007/OR=17.07) as major phenotypes (**Figure-2C-D**), indicating that the gnotobiotic mouse model is driving SCD. Notably, two instances of SD within the first week following cell injection (**Figure-1C**), indicates an unusually rapid onset of this condition.

Recent studies have highlighted that many bacterial metabolites and several gut bacterial species were depleted in patients with ischemic heart disease, a major cause of SCD ^8-10^. Host-microbiota interactions involving inflammatory and metabolic pathways have been linked to the pathogenesis of multiple immune-mediated diseases and metabolic conditions like diabetes and obesity^11-13^, also risk factors for cardiovascular diseases ^8-10^. The implications of our research extend beyond the scope of this study, prompting a thorough microscopic assessment of cardiac sub-micrometastasis and gut microbial assessment in SCD patients. While the lack of a cardiac tissue bank from SD patients poses a current obstacle, we are optimistic that forthcoming studies will overcome this hurdle, redefining our understanding of SCD. Nevertheless, there is a compelling need for further research on these enigmatic and highly lethal occult metastatic events.

In conclusion, our study introduces the first microbially preventable model of (SCD), providing novel insight on the pivotal role of the gut microbiota in regulating occult oncological events that contribute to such deaths. These findings underscore the need for further investigations to uncover the underlying microbes and their mechanisms involved in driving SCD.

## Methods

### Animal experiments

Three groups of C57BL/6NTac (B6) mice were included in the study: group 1, conventional (n=10); group 2, germ-free (n=22); and group 3, germ-free conventionalized (n=20) with a fecal sample from a C57BL/6NTac (B6) mouse donor at week 5 after birth. Up to 1 million Py230 mammary adenocarcinoma cells were injected into all three groups of mice at 8 weeks after birth. In most animals, injected cells were labeled with DNA barcode libraries as described previously^7^. The animals were monitored for tumor development or death for up to a year. At the study endpoint or in the event of death, primary tumors and/or vital organs were collected from the mice for further analysis. All procedures were reviewed and approved by the Mayo Clinic Institutional Animal Care and Use Committee.

### Microbial and clonal analysis

DNA was isolated from 10 mg of homogenized tumor, various tissues, or colonic fecal samples using the DNeasy Blood and Tissue Kit (Qiagen). Germ-free status was confirmed by PCR of the16s rRNA gene in colonic fecal DNA using a universal primer. For clonal analysis, 1 μg of genomic DNA from tumors and various tissues was spiked with ‘spike in’ controls and used as a template for PCR based NGS sequencing library preparation. The libraries were then sequenced on a HiSeq 4000. CIC calculator described elsewhere was used data filtering, thresholding, and calculating clonal output. For digital droplet PCR, 1 μg of genomic DNA was digested with EcoRI-HF enzyme (NEB) and analyzed using the QX100 system (Bio-Rad). QuantaSoft analysis software (Bio-Rad) was used to calculate the fractional abundance for each sample. The following primer sequences were used for amplification: forward–CCAGTAAGGCTGCTAGGAAAG, reverse–GATGCGAGTCAGAGAATGAGAG; probe sequence: -/56-FAM/TAGCCTTCTTAGGTGGCGTTGCAT/36-TAMSp/.

### Gene expression and dbGAP analysis

Total RNA was extracted from primary tumors using the TRI reagent and sequenced on the illumina Hiseq platform. Raw data was pre-processed and normalized using the DESeq2 package and rlog transformed. Differential gene expression analysis was conducted using the Weltch T test. Genes with a fold change > 1.5 or less than 0.7 and a p-value < 0.05 were considered differentially expressed (DE genes). The DE genes were fed for enrichment analysis with Enricher and checked for disease and drug associations. The obtained dbGAP genes were filtered based on an adjusted p-value < 0.05 and odds ratios were computed.

## Ethics statement

All procedures were reviewed and approved by Mayo Clinic Institutional Animal Care and Use Committee.

## Data availability statement

Except mouse tumor RNA-seq, all the data generated or analyzed during this study are included in this article. The RNA-seq datasets generated during the current study cannot be made publicly available due to intellectual property considerations. Sharing the data at this stage could compromise our ability to obtain necessary protections and may hinder future commercialization efforts. However, researchers interested in accessing the data for academic or collaborative purposes may contact corresponding author to discuss potential data sharing arrangements that comply with intellectual property and confidentiality obligations.

